# PGC-1β and ERRα promote glutamine metabolism and colorectal cancer survival via transcriptional regulation of PCK2

**DOI:** 10.1101/2022.05.20.492006

**Authors:** Danielle E. Frodyma, Thomas C. Troia, Chaitra Rao, Robert A. Svoboda, Jordan A. Berg, Dhananjay D. Shinde, Vinai C.Thomas, Robert E. Lewis, Kurt W. Fisher

## Abstract

Previous studies have shown that Peroxisome Proliferator-Activated Receptor Gamma, Coactivator 1 Beta (PGC-1β) and Estrogen-Related Receptor Alpha (ERRα) are over-expressed in colorectal cancer and promote tumor survival. In this study, we show that amino acid motif LRELL on PGC-1β is responsible for the physical interaction with ERRα and promotes ERRα mRNA and protein expression. We used RNAsequencing to determine the genes regulated by both PGC-1β & ERRα and found that mitochondrial Phosphoenolpyruvate Carboxykinase 2 (PCK2) was the gene that decreased most significantly after depletion of both genes. Depletion of PCK2 in colorectal cancer cells was sufficient to reduce anchorage-independent growth and inhibit glutamine utilization by the TCA cycle. Lastly, shRNA-mediated depletion of ERRα decreased anchorage-independent growth and glutamine metabolism, which could not be rescued by plasmid derived expression of PCK2. These findings suggest that transcriptional control of PCK2 is one mechanism used by PGC-1β and ERRα to promote glutamine metabolism and colorectal cancer cell survival.

## 1. Introduction

PGC-1 family proteins (PGC-1α, PGC-1β, and PPRC1) are transcriptional co-activators that bind a diverse array of transcription factors to promote the transcription of genes that regulate metabolism^1^. The combinations of PGC-1 family members and transcription factors are highly context dependent. We have previously shown that Peroxisome Proliferator-Activated Receptor Gamma, Coactivator 1 Beta (PGC-1β) and the transcription factor, Estrogen-Related Receptor Alpha (ERRα), are upregulated in colorectal cancer (CRC) in response to K-Ras mutations and that depletion of either protein decreases growth *in vitro* and *in vivo*^2,3^. However, the nature of the association between PGC-1β and ERRα and the genes they regulate in CRC has not been elucidated.

In this study, we used epitope tagging of endogenous PGC-1β to confirm the interaction between PGC-1β and ERRα. We then developed CRC cell lines with inducible expression of dual epitope-tagged PGC-1β and tested mutant PGC-1β proteins to identify the motif on PGC-1β required to bind ERRα. Next, we performed RNAsequencing to identify the genes that are regulated by both PGC-1β and ERRα and found that mitochondrial Phosphoenolpyruvate Carboxykinase 2 (PCK2) was the gene most significantly decreased after depletion of either protein. Using RNAi and liquid chromatography with tandem mass spectrometry, we explored the role of PCK2 in TCA cycle metabolism, glutamine utilization, and cell survival. Lastly, we generated a novel vector for stable over-expression of PCK2 to determine if PCK2 can rescue the loss of ERRα on L-glutamine utilization and anchorage-independent growth.

## 2. Materials and Methods

### Cell Culture

Colorectal cancer cell lines HCT116, T84, SW620, HT-29, SW480, and HCT15 were purchased from American Type Culture Collection (ATCC) and cultured in Dulbecco’s Modified Eagle’s Medium (DMEM) with 10% Fetal Bovine Serum (FBS), 2 mM L-glutamine and 1 mM sodium pyruvate at 37°C with ambient oxygen (O_2_) and 5% CO_2_.

### Lentiviral Transduction

For virus production, a 15-cm plate of HEK-293T cells at a confluence of 75% was transfected with 3 μg of pMD2.G, 6 μg of pPAX-2, and 12 μg of pLKO-shRNA-puro using 63 μL of polyethylenimine (PEI – 1 μg/μL; Polysciences - 24765) mixed in 750 μL of 10 mM HEPES pH 7.4 and 150 mM NaCl in water. The viral supernatant was cleared at 2,000 RPM for five minutes before being filtered through a 0.45 μM membrane filter then centrifuged at 12,000 RPM for two hours in a Sorval Lynx6000 with a F14 rotor. The resulting pellet was resuspended in four mL of media and 8 μL of polybrene (8 μg/μL) was added. 1 mL of the virus-polybrene solution was mixed with 1 mL of media containing 500,000 colorectal cancer cells and plated in one well of a six well plate.

DNA sequences are listed in Supplemental File 1.

### Epitope-tagging of endogenous PGC-1β

The homology directed repair (HDR) template for epitope tagging of endogenous PGC-1β was prepared by Gibson assembly of 3 pieces: 1) A 5’ prime homology arm containing approximately 750 base pairs of the genomic DNA upstream of the PGC-1β stop codon, a tobacco etch virus (TEV) cleavage site, and a twin strep 2 epitope tag; 2) a central region containing a triple FLAG epitope and P2A-neomycin resistance cassette acquired via restriction endonuclease digestion of pFETCH_Donor (Addgene: 63934); and 3) a 3’ prime homology arm containing approximately 750 base pairs of the 3’ untranslated region. Homology arms were ordered as gBlocks (Integrated DNA Technologies) and PCR-amplified. The three fragments were assembled with a Gibson Assembly Kit (New England Biolabs).

Then, HCT116 cells were transfected with the HDR template and pCAG-SpCas9-GFP-U6-gRNA (Addgene: 79144) expressing a gRNA targeted near the stop codon of PGC-1β. After two days, cells were moved to larger dishes, neomycin-selected, and clones were screened by PCR for correct genomic insertion.

DNA sequences are listed in Supplemental File 2.

### Immunoprecipitation of PGC-1β

Sample Preparation: Twenty 15-cm cell culture dishes of HCT116 or T84 cells expressing epitope tagged PGC-1β with 75% confluence were washed with PBS and each dish was lysed in 300 μL of RIPA lysis buffer (1% Triton X-100) with protease and phosphatase inhibitors (Halt Cocktail). Cells were sonicated and cleared in a centrifuge at 4°C for 20 minutes at 13,000 RPM in Thermo-Scientific Sorvall Lynx 6000 Centrifuge with F14 fixed angle rotor. Protein concentration was determined by BCA and sample were normalized to the same volume and concentration with additional RIPA buffer.

FLAG Immunoprecipitation: 200 μL of 50:50 magnetic FLAG bead slurry (Millipore-Simga; M8823) were washed twice with Tris Buffer Saline (TBS) and added to each sample and rotated overnight at 4°C. The beads were collected on a magnet and washed four times in 1 mL of TBS and eluted for three hours in 100 μL of 3X-FLAG peptide (100 ng/μL) in water.

Strep2 Immunoprecipitation: 100 μL of slurry of MagStrep “type3” XT beads (IBA biosciences, 2-4090-010) were washed in TBS and added to each sample and rotated overnight at 4°. The beads were collected on a magnet and washed four times in 1 mL of Tris Buffer Saline (TBS) and eluted for three hours in 100 μL of Buffer BXT (0.1 M Tris-Cl, 150 mM NaCl, 1 mM EDTA, 50 mM biotin, pH 8).

### Inducible PGC-1β Vector and Cell Line Generation

A full length human PGC-1β cDNA (Kind gift of Donald McDonnell, Duke) was PCR-amplified, digested, and ligated into AAVS1_Puro_Tet3G_3xFLAG_Twin_Strep (Addgene: 92099), and verified with bidirectional Sanger sequencing. Site directed mutagenesis was performed with the QuikChange Lightning Site-Directed Mutagenesis Kit (Agilent; 210513). Plasmids were integrated via dual transfection with pCAG-SpCas9-GFP-U6-gRNA (Addgene: 79144) expressing a gRNA targeted the AAVS1 T2 site.

DNA sequences for mutagenesis reactions are listed in Supplemental File 3.

### siRNA Transfections

The siRNA oligos (Dharmacon) targeting PGC-1β, ERRα, PCK2 or a non-targeting controls were used for targeted depletion of the colorectal cancer cells. For pooled transfections, two validated, individual ON-TARGET PLUS siRNAs were used at a final RNAi concentration of 40nM and was added to 5μL of RNAiMAX (ThermoFisher, 13778150) and 500μL Hank Buffer Salt Solution without sodium bicarb. The mixture was added to 300,000 cells in 1.5-2mL of media without antibiotics in a 6-well plate. All transfections were conducted for 72-hours before analysis.

RNA sequences for transfections are listed in Supplemental File 4.

### RNA Sequencing and Analysis

RNA sequencing (RNA-seq) analysis was conducted by the UNMC Genomics Core. Cells were harvested using 0.5 mL TRIzol (ThermoFisher Scientific) and stored at -80°C until RNA extraction was performed. RNA was extracted using RNeasy spin columns (Qiagen) per manufacturer’s protocol. Final RNA was eluted with nuclease-free water and quantified using the NanoDrop 2000 (ThermoFisher Scientific). Three biological replicates of non-targeting control, PGC-1*β*, or ERR*α* knockdown were completed using two separate siRNA oligos for each condition. Unstranded (poly A only) RNA sequencing libraries and 500 ng of total RNA for each of the samples were prepared per manufacturer’s suggested protocol using the TrueSeq mRNA Protocol Kit (Illumina). Purified libraries were pooled at a 0.9 pM concentration and sequenced on an Illumina NextSeq550 instrument and 75 bp paired end sequencing was performed. Libraries were normalized and equal volumes were pooled in preparation for sequence analysis. Raw sequence data has been deposited as GSE147905 in the National Center for Biotechnology Information Gene Expression Omnibus. Sequence reads were preprocessed using XPRESSpipe(v0.4.1)^4^, with adapter sequences AGATCGGAAGAGCACACGTCTGAACTCCAGTCA and AGATCGGAAGAGCGTCGTGTAGGGAAAGAGTGT. Reads were processed using H. sapiens GRCh38.13 Ensembl release 99. Differential expression analysis was performed using XPRESSpipe wrapper for DESeq2 (v1.22.1)^5^. Differentially expressed genes were further visualized using XPRESSplot. Isoform abundance analysis was performed using XPRESSpipe wrapper for Cufflinks (v2.1.1)^6^ and IGV (v2.4.19)^7^. Scripts used to perform these analyses can be found at https://github.com/j-berg/frodyma_2020.

### Western Blot Analyses

Whole cell lysate extracts were prepared in radioimmunoprecipitation assay (RIPA) buffer that was comprised of 50 mM Tris-HCl, 1% NP-40, 0.5% sodium deoxycholate, 0.1% sodium dodecyl sulfate, 150 mM NaCl, 2 mM EDTA, 2 mM EGTA, and addition of a protease and phosphatase inhibitor cocktail (Halt, ThermoFisher Scientific). A BCA protein assay (Promega) was used to determine protein concentration. An 8% Acrylamide SDS-PAGE was used to separate out the protein and nitrocellulose membranes were blocked in Odyssey TBS blocking buffer (LI-COR Biosciences) for at least 30 minutes at room temperature. The primary antibody was allowed to hybridize at least overnight at 4°C. The PCK2 (8565) antibody was obtained from Cell Signaling Technologies and used at a concentration of 1:2000. The PGC-1β (NBP1-28722) antibody was purchased from NovusBio and used at a concentration of 1:1000. The ERRα (ab76228) antibody was purchased from Abcam and used at a concentration of 1:1000. The FLAG epitope (F1804) antibody was purchased from Millipore-Sigma and used at a concentration of 1:5000. The Strep2 epitope (Ab02208-1.1) antibody was purchased from Absolute Antibody and used at a concentration of 1:2000. The β-actin (sc-47778) and α-tubulin (sc-5286) antibodies were purchased from Santa Cruz Biotechnology and used at a concentration of 1:2000. The anti-ALFA recombinant nanobody-rabbit Fc fusion (N1583) was obtained for NanoTag Biotechnologies and used at a concentration of 1:2000. IRDye 800CW and 680RD secondary antibodies (LI-COR Biosciences) were diluted 1:10,000 in 0.1% TBS-Tween and imaged on an Odyssey Scanner (LI-COR Biosciences).

### L-Glutamine Utilization Assays

The Seahorse XFe96 Metabolic Flux Analyzer (Agilent) was used to measure Oxygen Consumption Rate (OCR) in the presence of only 2 mM L-glutamine as a substrate. The day before the experiment, the FluxPak plates were hydrated in water and incubated at 37°C with ambient CO2. The afternoon before the assay, 40,000 cells were plated in each well of a 96 well assay plate in 12 replicates in regular media. On the day of the experiment, the media was removed and the cells were washed twice with 1 mL of PBS then covered in 180μL of XF DMEM medium pH 7.4, (Agilent 103575-100) with 2mM glutamine (Agilent 103579-100). The cells were incubated in this media at 37°C with ambient CO_2_ for 1 hour prior to beginning the experiment. The water was removed from the FluxPak and calibration media was added and incubated at 37°C with ambient atmosphere for an hour prior to the experiment. Basal OCR was measured four times for three minutes with mixing between measurements to ensure stability and the last measured was used for statistical evaluation.

### Intracellular Metabolite Analysis

HCT116 cells were transfected in five biological replicates, as previously described. 72-hours after transfection, the cells were harvested and counted. After washing in saline solution, the cell pellet was resuspended into 1 mL of ice-cold 2:2:1 MeOH: ACN: H2O (v/v/v) containing 10 μM stable isotope-labeled canonical amino acid mix (Cambridge Isotope Laboratories, Inc) as internal standards. The cells were subsequently lysed in a reciprocal shaker with 0.1 mm glass beads and the samples were centrifuged for 15 minutes at 13,000 rpm at 4°C. The supernatant was removed and evaporated to dryness in the SpeedVac. The samples were reconstituted in 100 μL of resuspension buffer containing 20% ACN and 10 mM ammonium acetate, before LC-MS/MS analysis.

Chromatographic separation and mass Spectrometry detection were performed using a Shimadzu Nexera ultra-high-performance liquid chromatography (UHPLC) and triple-quadrupole-ion trap hybrid Mass spectrometer (QTRAP 6500 from Sciex, USA), equipped with an ESI source. The chromatographic separation of metabolites was achieved on a XSelect (150 × 2.0 mm id; particle size 1.7 μm) analytical column maintained at 40°C. The optimum mobile phase consisted of 10 mM tributylamine with 5 mM acetic acid in LC-MS grade water containing 2% isopropanol as buffer A and isopropanol as solvent B. The gradient elution performed as: time zero to five min, 0% solvent B; next 4 min, 2% solvent B; 0.5 min, 6% solvent B; 2 min, 6% solvent B; for next 0.5 min, solvent B was increased to 11% and maintained for 1.5 min; at 35 min, solvent B was increased to 28% for next 2 min and then to 53% in 1 min and maintained for next 6.5 min. Solvent B was reduced to 0% and maintained to equilibrate column till the next injection. The flow rate was 0.4 mL/min, and the total run time was 33 min, and the autosampler temperature was 10°C. The data acquisition was under the control of MultiQuant software (Sciex, USA). The mass spectrometer was operated in positive as well as negative ion mode using polarity switching. Ions were acquired in multiple reaction monitoring (MRM) mode. MRM details for the selected metabolites were as follows: Oxaloacetate, 131.0/ 87.0; Phosphoenolpyruvate, 167.0/ 79.0; Citrate/Isocitrate pool, 191.0/ 111.0; Fumarate, 115.0/ 71.0; Succinate, 117.0/ 99.0. The retention time of each metabolite was confirmed by the 13C-labelled yeast metabolite extract, which was used as the qualitative standard (Cambridge Isotope Laboratories, Inc). Optimized spray voltage was at 5.5 kV for positive and 4.2 kV for negative mode, ESI source temperature was at 400°C, nitrogen was used as curtain gas, gas 1 and gas 2 at pressure 30, 40 and 40 arbitrary units, respectively. Declustering potential in positive and negative mode was optimized at 65 and -65 volts.

### PCK2-ALFA Plasmid for Stable Expression

A full length PCK2 cDNA was obtained through Addgene (plasmid: 23715) and was PCR amplified with a 3’ primer containing a single ALFA tag. The resulting PCR product was digested and ligated into pcDNA-hEF-1α-neomycin resistance. The final product was verified with Sanger sequencing from both directions and incorporated into cells using PEI transfection and selection with G418 (InvivoGen, ant-gn-5).

### Confocal Microscopy

200,000 cells with PCK2-ALFA expression were plated in 6-well plates containing 2 glass cover slips (12 mm; Deckglaser) in DMEM medium with 10% FBS. The following day, they were stained with 100 nM MitoTracker Deep Red (Invitrogen) for 30 minutes before washing and formalin fixation. Cells were then stained with a 1:3,000 dilution of FluoTag®-X2 anti-ALFA conjugated to Atto-488 (NanoTag Biotechnologies; N1502-At488) per the manufacturer’s directions. Cells were mounted to glass slides using Fluoromeount G DAPI (SouthernBiotech; 0100-20) and imaged on a Zeiss 800 CLSM with Airyscan at the UNMC Advanced Microscopy Core Facility.

### Statistical Analysis

P values were calculated using Prism Software (GraphPad, v8.4.2, La Jolla, CA). A P value of less than 0.05 was considered statistically significant. The statistical significance of these results was evaluated using one way ANOVA with multiple comparisons to knockdown in each cell line. The cell metabolic capacity assays were statistically evaluated using an unpaired, two-sided t-test to compare the effects of PCK2 depletion to control cells. Data are shown as mean +/- standard deviation (SD) unless otherwise noted.

## 3. Results

### 3.1 Endogenous PGC-1β interacts with ERRα, promotes ERRα expression, and anchorage-independent growth

We have previously shown that shRNA-mediated depletion of PGC-1β caused ERRα protein levels to decrease in human CRC cell line, HCT116^2^. Here, we determined how robust this observation is by using lentiviral-mediated delivery of shRNAs to decrease PGC-1β expression in a panel of human CRC cell lines and immunoblotted for PGC-1β, ERRα, and measured anchorage-independent growth by colony formation in soft agar. Depletion of PGC-1β caused a decrease in ERRα protein levels and anchorage independent growth in a panel of K-Ras mutant CRC cell lines (Fig. 1A-B).

**Figure 1.**
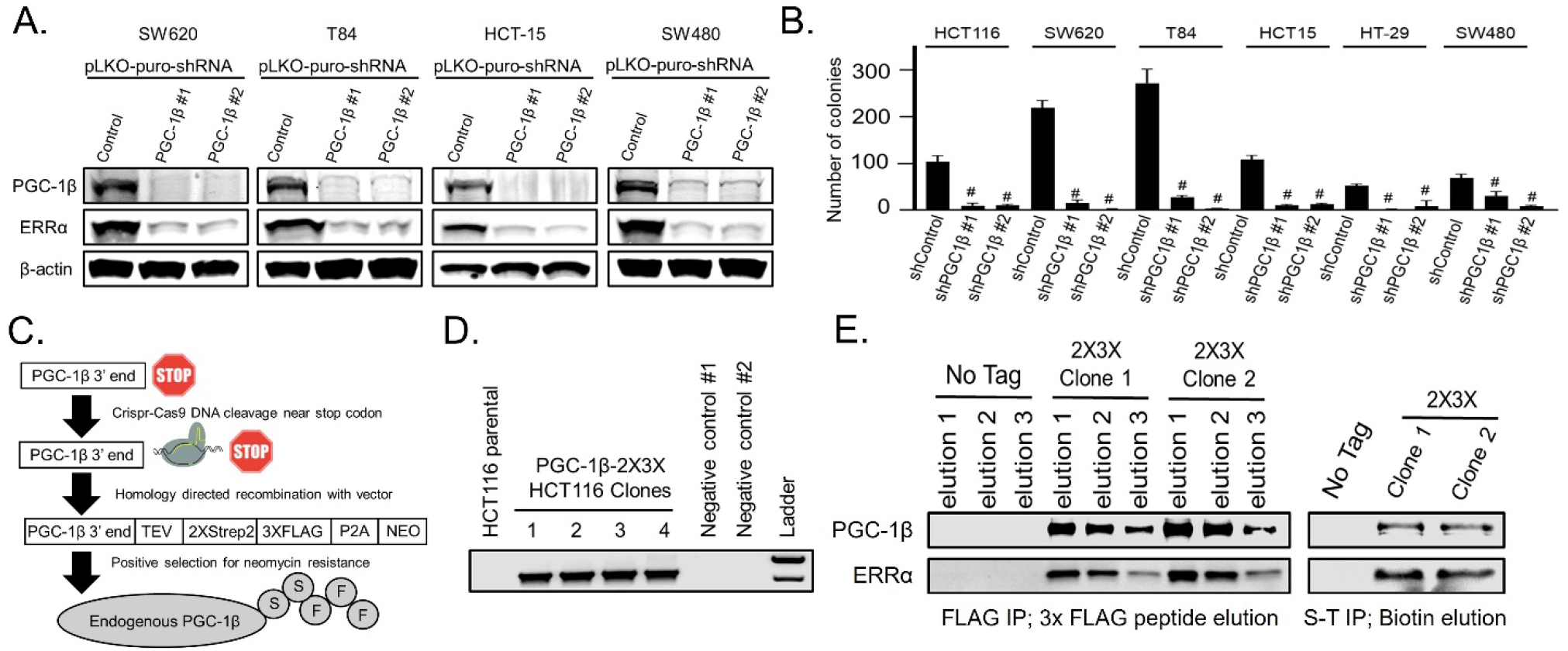
Endogenous PGC-1β interacts with ERRα, promotes ERRα expression, and increases anchorage-independent growth. A) Lentiviral mediated transduction was used to express shRNAs for either a non-targeting control or two independent sequences targeting PGC-1β. The cells were lysed after 72 hours and immunoblotted for PkfisGC-1β, ERRα, and loading control β-actin. B) Colony formation in soft agar was measured after two weeks in cells infected with the same shRNAs from panel A. C) Illustrative representation of the strategy used to knock in twin Strep2 and triple FLAG epitope tags onto the C-terminus of endogenous PGC-1β using CRIRSP-Cas9 mediated genome cutting with homology directed repair and neomycin selection. D) Agarose gel showing PCR verification of correct genomic insertion of the epitope tags from part C in four clones of HCT116 cells. E) HCT116 cells with epitope tagged endogenous PGC-1β were immunoprecipitated for either the FLAG or the Strep2 epitope and eluted with either the 3X-FLAG peptide or 50 mM biotin. Eluates were immunoblotted for PGC-1β and ERRα. # = p-value less than 0.001 when compared to the non-targeting control using one way ANOVA with multiple comparisons. S-T IP= Strep-Tactin immunoprecipitation.

PGC-1 proteins are known to bind transcription factors, but the physical interaction between the PGC-1β and ERRα has not been explored in detail. To investigate the physical interaction between PGC-1β and ERRα, we generated a vector for epitope-tagging of endogenous PGC-1β using homology directed repair (HDR). Using CRISPR-Cas9, we generated a double stranded break adjacent to the stop codon of PGC-1β and used our plasmid as a template for HDR to eliminate the stop codon and incorporate twin Strep2 triple FLAG epitopes and a neomycin resistance cassette (Fig 1C). After neomycin selection, clones were screened by PCR to confirm the correct genomic insertion (Fig. 1D). Endogenous PGC-1β was immunoprecipitated by its FLAG epitopes and eluted with the 3X-FLAG peptide or immunoprecipitated by the Strep2 epitopes and eluted with the biotin. Immunoblotting of the eluates showed both PGC-1β and ERRα, confirming their interaction (Fig. 1E). These findings suggest PGC-1β binds ERRα to promote ERRα transcriptional activity.

### 3.2 PGC-1β requires its LRELL motif at amino acids 343-347 to interact with ERRα

PGC-1 family proteins have been shown to use LxxLL amino acid motifs to bind transcription factors^8–11^. To determine the motif(s) required by PGC-1β to bind ERRα, we developed cell lines with inducible expression of N-terminus twin Strep2 triple FLAG epitope-tagged PGC-1β under doxycycline inducible expression from the AAVS1 safe harbor locus and found the increased levels of PGC-1β also causes a modest induction of ERRα protein levels (Fig. 2A). We then made a series of PGC-1β mutant proteins to assess the role of each LxxLL motif in binding ERRα. To be in accordance with previous labeling from the literature^8^ we maintained the same labeling system: Motif 1 LLAEL (amino acids 92-96), Motif 2 LKQLL (amino acids 156-160), Motif 3 LRELL (amino acids 343-347), and Motif 4 LLSHL (amino acids 664-668). Technically, motifs 1 and 4 are reversed but we wanted to directly assess their role in ERRα binding since there is literature evidence to suggest these motifs may be functional in other PGC-1 family members^12^. Motifs were inactivated by mutating all leucines to alanines (LxxLL → AxxAA or LLxxL → AAxxA). Using this strategy, we created one quadruple PGC-1β mutant where all four motifs were inactivated (zero LxxLL motifs) and four triple mutants where only one LxxLL motif was left non-mutated (Only LxxLL motif #1, Only LxxLL motif #2, Only LxxLL motif #3, and Only LxxLL motif #4). The five mutant and wild type PGC-1β cDNAs were integrated into the AAVS1 safe harbor locus of HCT116 cells and the expressed proteins immunoprecipitated using the FLAG or Strep2 epitopes in separate experiments. The eluates were immunoblotted for PGC-1β and ERRα and showed that the mutant PGC-1β that was functional at only the LxxLL motif #3 (LRELL) immunoprecipitated the same amount as wild type PGC-1β (Fig. 2B). To test the role of the LRELL motif, we made two additional mutants that had all three leucines mutated to alanines (LRELL → AREAA mutant) or the RE mutated to alanines (LRELL → LAALL mutant). The PGC-1β AREAA mutant cDNA was integrated into both HCT116 and T84 cells and tested against wild type PGC-1β. Loss of the leucines in motif #3 eliminated ERRα binding in both cell lines (Fig. 2C). To test the role of amino acids RE in motif #3 on ERRα binding we generated a PGC-1β LAALL mutant cDNA that was integrated into HCT116 cells and tested against wild type PGC-1β. Mutation of amino acids RE showed only a partial loss of ERRα binding (Fig 2D). Lastly, we tested if the PGC-1β mutant that can’t bind ERRα could induce ERRα expression and found that the PGC-1β AREAA mutant was unable to induce ERRα expression compared to wild type PGC-1β (Fig. 2E). Overall, these findings suggest that all five amino acids of the PGC-1β LRELL motif are required for optimal ERRα protein binding, which causes increased levels of ERRα protein.

**Figure 2.**
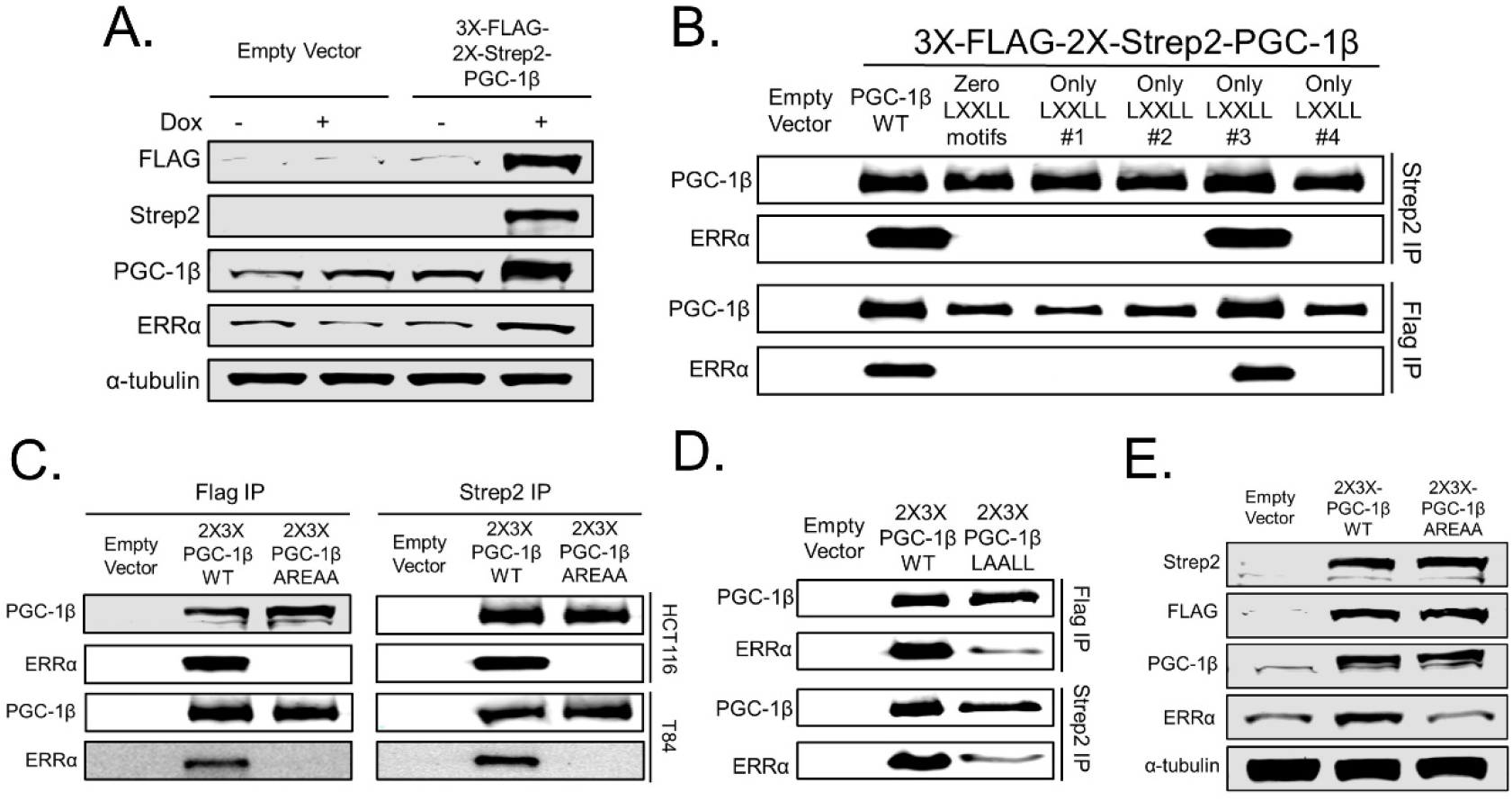
PGC-1β requires the LRELL motif at amino acids 343-347 to interact with ERRα. A) HCT116 cells were cut at the AAVS1 safe harbor locus with CRISPR-Cas9 and homology-directed repair and puromycin selection were used to integrate an inducible full length PGC-1β cDNA with triple FLAG and twin Strep2 epitope tags on the N-terminus. Immunoblots for PGC-1β, Strep2, and FLAG confirm tagged protein induction, which causes increased ERRα expression. B) HCT116 cells with either an empty vector, wild type dual epitope tagged PGC-1β, or five different PGC-1β mutants were immunoprecipitated for either the FLAG epitope or the Strep2 epitope and eluted with a triple FLAG peptide or 50 mM biotin, respectively. Eluates were subjected to immunoblot for PGC-1β and ERRα. C) HCT116 and T84 cells with either an empty vector, wild type dual epitope tagged PGC-1β, or PGC-1β mutated at the LRELL motif to AREAA were immunoprecipitated as before and immunoblotted for PGC-1β and ERRα. D) HCT116 cells with either an empty vector, wild type dual epitope tagged PGC-1β, or PGC-1β mutated at the LRELL motif to LAALL were immunoprecipitated as before and immunoblotted for PGC-1β and ERRα. E) HCT116 cells expressing either wild type or the AREAA mutant PGC-1β from the AAVS1 safe harbor locus were treated with doxycycline for 24 hours and lysed for western blot to assess ERRα induction.

### 3.3 PGC-1β and ERRα promote PCK2 expression

The genes regulated by PGC-1β and ERRα have only been examined in detail in breast cancer and normal liver tissue^8,13,14^, but not in CRC. To determine which genes were regulated by PGC-1β and ERRα in CRC, we validated two siRNAs that targeted either protein (Fig. 3A) and that loss of PGC-1β caused decreased levels of ERRα, but decreased levels of ERRα did not alter PGC-1β expression. To determine the genes regulated by PGC-1β and ERRα, we transfected HCT116 cells with siRNAs targeting either PGC-1β, ERRα, or non-targeting controls and collected total RNA after 72 hours for RNA sequencing (Supplemental File 5). First, depletion of PGC-1β led to a dramatic decrease in ERRα mRNA, consistent with the model in the literature^13,15^ that binding of PGC-1β to ERRα promotes ERRα binding to its own promoter to increase ERRα mRNA transcription (Fig. 3B). Secondly, mitochondrial Phosphoenolpyruvate Carboxykinase 2 (PCK2) decreased the most after depletion of both PGC-1β and ERRα (Figs.

**Figure 3.**
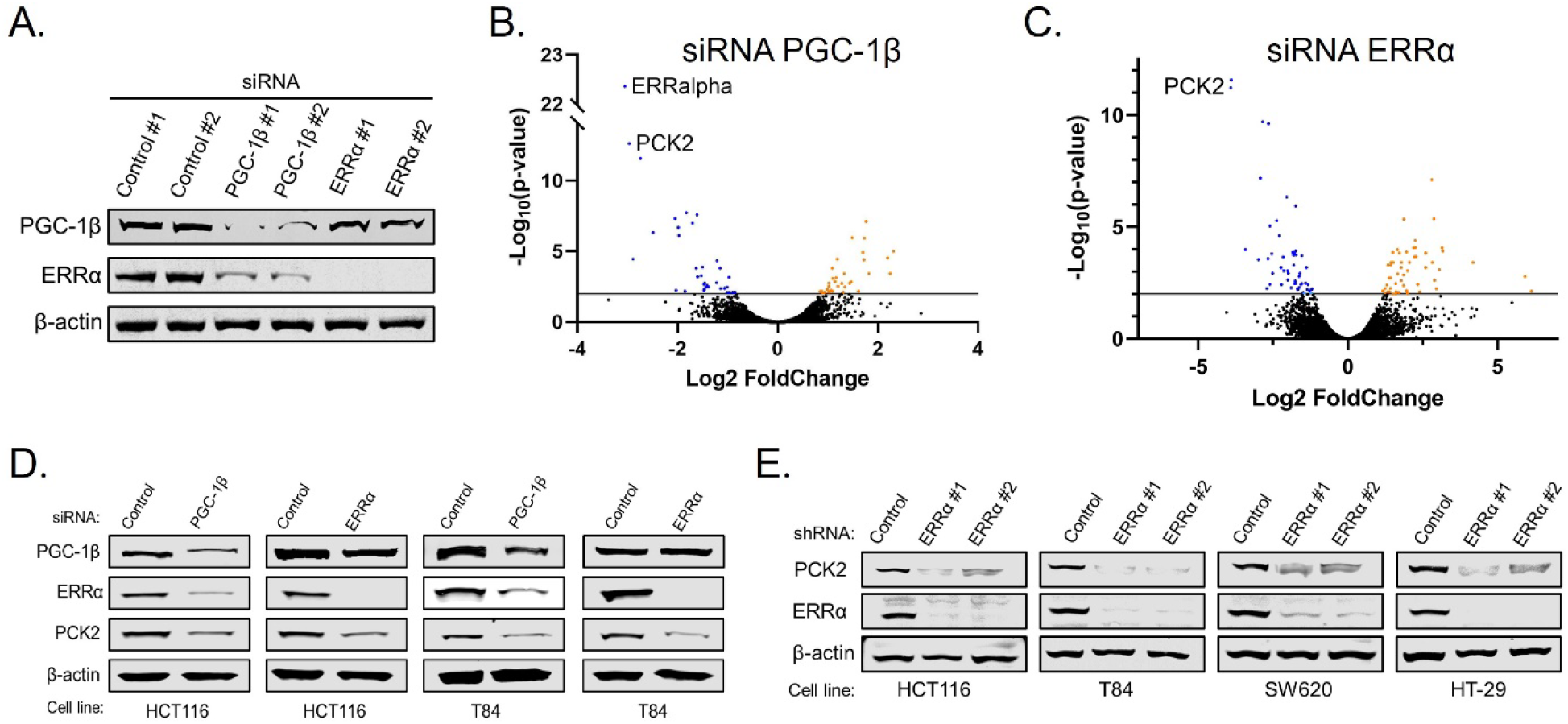
PGC-1β and ERRα promote PCK2 expression. A) HCT116 cells were transfected with siRNAs targeting either PGC-1β or ERRα or a non-targeting control. After 72 hours, cells were lysed and immunoblotted to confirm the depletion of the target proteins. B-C) Volcano plots of RNAsequncing data from total RNA analysis 72 hours after knockdown of PGC-1β or ERRα compared to a non-targeting control. D) Colorectal cancer cell lines were transfected with siRNAs targeting either PGC-1β or ERRα and after 72 hours were immunoblotted for PCK2. E) Colorectal cancer cell lines were transduced with lentiviruses expressing shRNAs targeting ERRα or a non-targeting control and after 96 hours were lysed and immunoblotted for PCK2 and ERRα.

3B-C). PCK2 is localized to the mitochondria and catalyzes the irreversible conversion of oxaloacetate (OAA) to phosphoenolpyruvate (PEP). Lastly, several other genes that regulate amino acid metabolism decreased after depletion of both PGC-1β and ERRα, suggesting that these two proteins cooperate to promote amino acid incorporation and metabolism to increase survival of colorectal cancer cells.

To confirm that the mRNA changes seen after depletion of PGC-1β and ERRα lead to changes in PCK2 protein expression we performed transient knockdown of either protein in two colorectal cancer cell lines and observed decreased levels of PCK2 by Western blot (Figs. 3D). Lastly, we used lentiviral mediated delivery of shRNA targeting ERRα to reduce ERRα expression in three CRC cell lines and observed decreased levels of PCK2 by Western blot (Figs. 3E). These findings suggest the transcriptional control of PCK2 levels by PGC-1β and ERRα is a mechanism to control amino acid metabolism in CRC.

### 3.4 PCK2 promotes anchorage independent growth and glutamine utilization

PCK2 mRNA has been shown to be upregulated by mutant K-Ras^16^, but its functional role has not been explored in CRC. We have previous shown that PGC-1β expression is dependent on mutant K-Ras signaling^2^, suggesting that transcriptional control of PCK2 by PGC-1β is part of oncogenic K-Ras mediated metabolic reprogramming. To assess the role of PCK2 in CRC survival we used lentiviral-mediated delivery of shRNAs targeting PCK2 to deplete PCK2 protein (Fig. 4A) and performed anchorage-independent growth studies. We found that depletion of PCK2 caused a significant decrease in soft agar colony formation in a panel of K-Ras mutated CRC cell lines (Figs. 4B-C).

**Figure 4.**
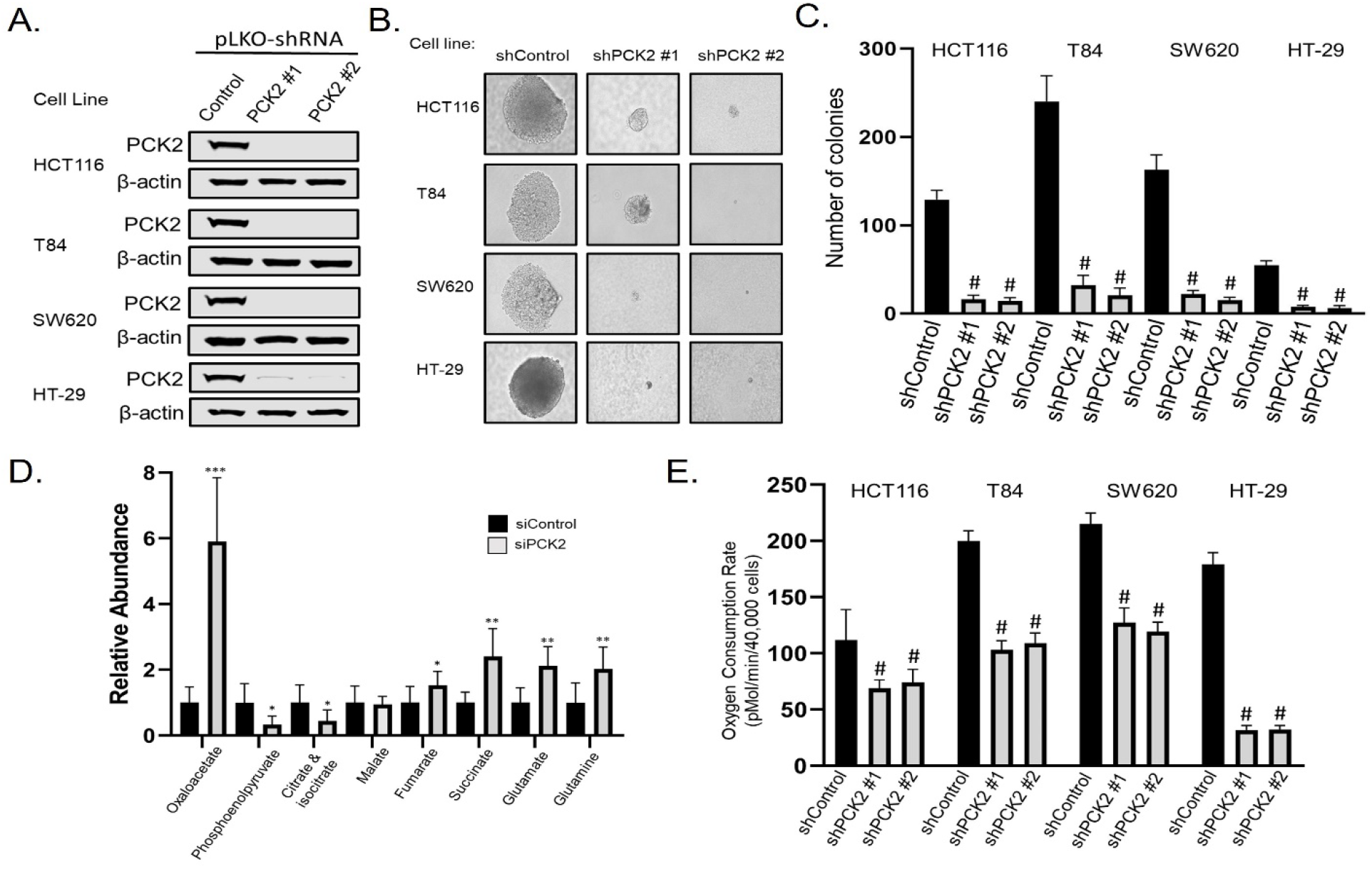
PCK2 promotes anchorage independent growth and glutamine utilization. A) Colorectal cancer cell lines were transduced with lentivirus producing shRNAs targeting PCK2 and subjected to immunoblot after 72 hours to confirm knockdown. B) Representative photomicrographs of colonies in soft agar two weeks after shRNA mediated depletion of PCK2 in four CRC cell lines C) Quantification of colonies from panel B. D) Intracellular metabolites were measured by LC-MS/MS 72 hours after siRNA-mediated depletion of PCK2. E) Oxygen consumption rates were measured using a Seahorse XFe96 analyzer in colorectal cancer cell lines 72 hours after shRNA-mediated knockdown of PCK2 using only L-glutamine as a substrate. * = p-value less than 0.05; ** = p-value less than 0.01, *** = p-value less than 0.001. # = p-value less than 0.001 when compared to the non-targeting control using one way ANOVA with multiple comparisons.

To determine how depletion of PCK2 alters metabolism, we transiently depleted PCK2 in HCT116 cells and measured intracellular metabolites using liquid chromatography and tandem mass spectrometry (Fig. 4D). PCK2 is responsible for the irreversible conversion of oxaloacetate (OAA) to phosphoenolpyruvate (PEP) within the mitochondria. As expected, depletion of PCK2 caused a six-fold increase in total oxaloacetate (OAA) levels and a 55% decrease in total phosphoenolpyruvate (PEP) levels. Unexpectedly, elevated levels of OAA did not appear to be converted into citrate and isocitrate, as the pool of citrate/isocitrate was lower in cells with PCK2 depletion. Depletion of PCK2 also caused an accumulation of upstream TCA metabolites fumarate, succinate, glutamate, and glutamine, suggesting that glutamine flux through the TCA cycle was diminished in the absence of PCK2 activity. To assess L-glutamine utilization, we used shRNAs to deplete PCK2 in a panel of CRC cell lines and measured their oxygen consumption rate using a Seahorse Metabolic Analyzer using only L-glutamine as a substrate. Depletion of PCK2 caused a significant decrease in L-glutamine oxidation in four CRC cell lines (Fig. 4E). Overall, these findings suggest that PCK2 can control glutamine flux through the TCA cycle to promote CRC cell survival.

### 3.5 ERRα depletion causes loss of anchorage-independent growth and glutamine utilization that is not rescued by over-expression of PCK2

To assess if PCK2 can rescue the loss of ERRα activity, we developed a novel plasmid for stable PCK2 expression. PCK2 is anchored to the mitochondrial membrane at its N-terminus so we added an ALFA epitope tag^17^ to the C-terminus and expressed it via a human EF-1α promoter in two CRC cell lines after neomycin selection (Fig. 5A). We confirmed that the PCK2-ALFA protein was successfully localized to the mitochondria by performing direct immunofluorescence for the ALFA epitope using an ALFA recognizing nanobody conjugated to Atto-488 and compared it MitoTracker Far Red staining (Fig. 5B).

**Figure 5.**
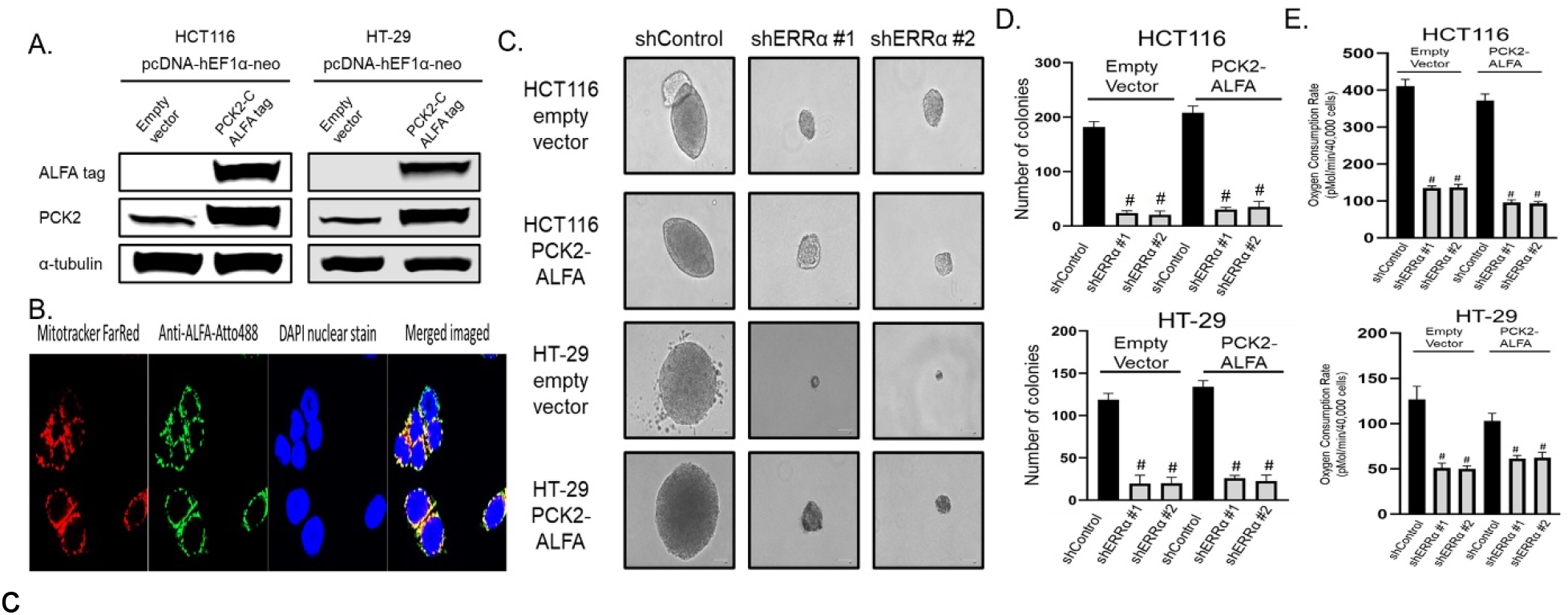
ERRα depletion causes loss of anchorage-independent growth and glutamine utilization and that is not rescued by elevated PCK2 levels. A) Immunoblot of colorectal cancer cell lines showing stable expression of PCK2 with a C-terminus ALFA epitope tag after neomycin selection. B) Confocal photomicrographs showing the overlap of PCK2 with mitochondrial dye. C) Representative photomicrographs of colonies in soft agar two weeks after shRNA mediated depletion of ERRα in two CRC cell lines with plasmid derived PCK2 expression or an empty vector control D) Quantification of colonies from panel C. D) Oxygen consumption rates were measured using a Seahorse XFe96 analyzer in colorectal cancer cell lines from panels C and D using only L-glutamine as a substrate. # = p-value less than 0.001 when compared to the non-targeting control using one way ANOVA with multiple comparisons.

To determine if over-expression of PCK2 could rescue the loss of ERRα, we used lentiviral mediated delivery of shRNAs targeting ERRα in two cell lines stably expressing PCK2-ALFA or an empty vector and performed anchorage-independent growth and L-glutamine utilization assays. Loss of ERRα caused a significant loss of colony formation and L-glutamine metabolism that was not rescued by the over-expression of PCK2 (Fig. 5C-E). These findings suggest that transcriptional control of PCK2 expression is one mechanism used PGC-1β and ERRα to promote glutamine metabolism and CRC cell survival, but that other PGC-1β and ERRα target genes are also important to CRC metabolism and survival.

## 4. Discussion

Here we show that PGC-1β and ERRα physically interact and promote genes that promote amino acid metabolism. PGC-1β has been shown to regulate several metabolic processes in other model systems^14,18–20^, but the regulation of PCK2 and amino acid metabolism is a novel observation in CRC that differs from previous studies. Our study is the first to examine the role of PCK2 in CRC metabolism and survival and our observations suggest that PCK2 maximizes flux through the TCA cycle by converting OAA to PEP to promote cell survival. Our finding that elevated levels of OAA after siRNA-mediated depletion of PCK2 were not converted to citrate and isocitrate suggests that there is insufficient levels of either citrate synthetase or its other substrate Acetyl-CoA. Using Western blot, we were able to easily detect citrate synthetase in all CRC cell lines tested (data not shown). We did not measure levels of Acetyl-CoA in the mitochondria. These data are consistent with reports that the mitochondrial pyruvate complex is down-regulated in colorectal cancer^21,22^, which would limit the import of pyruvate into the mitochondria for conversion into Acetyl-CoA. These findings suggest intramitochondrial levels of Acetyl-CoA are insufficient to convert excess OAA to citrate and that conversion of OAA to PEP, presumably for mitochondrial export, is the most efficient method for utilizing L-glutamine by the TCA cycle in CRC.

Our data show that PGC-1β uses all five amino acids of its LRELL motif to bind ERRα protein, but no other LxxLL or LLxxL motif appears to bind ERRα, which suggests to us that PGC-1β can only bind one ERRα molecule at a time. ERRα has over 800,000 predicted bindings sites in the human genome, raising the possibility that additional transcription factors are required for gene selection by PGC-1β. Here, we have created several novel PGC-1β mutant proteins that can be used to assess the role of each LxxLL and LLxxL motif alone or in combination to determine their role in transcription factor binding and to discover new components of PGC-1β signaling. Additionally, PGC-1 family proteins have been shown to bind Host Cell Factor proteins through a DHDY motif^23^, which represents another potential mechanism that PGC-1β may use for gene selection. Host Cell Factor 1 and 2 have been proposed to bridge transcriptional co-activators, such as PGC-1 family proteins, to DNA binding via transcription factor binding at their N-terminus Kelch repeat domains, but this process has not been fully explored.

Lastly, our study exclusively involves colorectal cancer cell lines with K-Ras mutations. We have previously shown that PGC-1β expression is dependent on mutant K-Ras^2,3^, but we have not investigated whether these findings extend to different genetic background, such as B-Raf mutant tumors, HER-2 over-expressing tumors, or tumors that are wild type for these mutations. Further work to understand how these genetic background affect PGC-1β-ERRα signaling would be beneficial before attempting to target their signaling or PCK2 activity as a therapeutic strategy.

## 5. Conclusion

Inhibiting amino acid metabolism in CRC with K-Ras mutations by targeting PGC-1β signaling has the potential to provide cancer cell specific therapy without subjecting the patient to the stresses of global metabolic restriction.

## Author Contributions

Conceptualization, D.F., K.F., and R.L.; data curation, D.F., T.T., J.A.B., D.S., V.T., R.S., and C.R.; writing—original draft preparation, D.F.; writing—review and editing, D.F., C.R., J.A.B, K.F., and R.L.; visualization, D.F.; supervision, K.W., and R.L.; funding acquisition, R.L. and K.F. All authors have read and agreed to the published version of the manuscript.

## References

1. Hock MB, Kralli A. Transcriptional control of mitochondrial biogenesis and function. Annu Rev Physiol. 2009;71:177–203. PMID: 19575678

2. Fisher KW, Das B, Kim HS, Clymer BK, Gehring D, Smith DR, Costanzo-Garvey DL, Fernandez MR, Brattain MG, Kelly DL, MacMillan J, White MA, Lewis RE. AMPK Promotes Aberrant PGC1beta Expression To Support Human Colon Tumor Cell Survival. Mol Cell Biol. 2015 Nov;35(22):3866–79.

3. McCall JL, Gehring D, Clymer BK, Fisher KW, Das B, Kelly DL, Kim H, White MA, Lewis RE. KSR1 and EPHB4 Regulate Myc and PGC1beta To Promote Survival of Human Colon Tumors. Mol Cell Biol. 2016 Sep 1;36(17):2246–61.

4. Berg JA, Belyeu JR, Morgan JT, Ouyang Y, Bott AJ, Quinlan AR, Gertz J, Rutter J. XPRESSyourself: Enhancing, standardizing, and automating ribosome profiling computational analyses yields improved insight into data. PLOS Comput Biol. Public Library of Science; 2020 Jan 31;16(1):e1007625.

5. Love MI, Huber W, Anders S. Moderated estimation of fold change and dispersion for RNA-seq data with DESeq2. Genome Biol. 2014;15(12):550. PMCID: PMC4302049

6. C T, A R, L G, G P, D K, Dr K, H P, Sl S, Jl R, L P. Differential gene and transcript expression analysis of RNA-seq experiments with TopHat and Cufflinks. Nat Protoc [Internet]. Nat Protoc; 2012 Mar 1 [cited 2022 Jun 14];7(3). Available from: https://pubmed.ncbi.nlm.nih.gov/22383036/ PMID: 22383036

7. Robinson JT, Thorvaldsdóttir H, Winckler W, Guttman M, Lander ES, Getz G, Mesirov JP. Integrative genomics viewer. Nat Biotechnol. 2011 Jan;29(1):24–26. PMCID: PMC3346182

8. Chang C yi, Kazmin D, Jasper JS, Kunder R, Zuercher WJ, McDonnell DP. The metabolic regulator ERRα, a downstream target of HER2/IGF-1R, as a therapeutic target in breast cancer. Cancer Cell. 2011 Oct 18;20(4):500–510. PMCID: PMC3199323

9. Park S, Chang C yi, Safi R, Liu X, Baldi R, Jasper JS, Anderson GR, Liu T, Rathmell JC, Dewhirst MW, Wood KC, Locasale JW, McDonnell DP. ERRα Regulated Lactate Metabolism Contributes to Resistance to Targeted Therapies in Breast Cancer. Cell Rep. 2016 Apr 12;15(2):323–335. PMCID: PMC4833658

10. Rha GB, Wu G, Shoelson SE, Chi YI. Multiple binding modes between HNF4alpha and the LXXLL motifs of PGC-1alpha lead to full activation. J Biol Chem. 2009 Dec 11;284(50):35165–35176. PMCID: PMC2787377

11. Delerive P, Wu Y, Burris TP, Chin WW, Suen CS. PGC-1 functions as a transcriptional coactivator for the retinoid X receptors. J Biol Chem. 2002 Feb 8;277(6):3913–3917. PMID: 11714715

12. Kressler D, Schreiber SN, Knutti D, Kralli A. The PGC-1-related protein PERC is a selective coactivator of estrogen receptor alpha. J Biol Chem. 2002 Apr 19;277(16):13918–13925. PMID: 11854298

13. Deblois G, Hall JA, Perry MC, Laganière J, Ghahremani M, Park M, Hallett M, Giguère V. Genome-wide identification of direct target genes implicates estrogen-related receptor alpha as a determinant of breast cancer heterogeneity. Cancer Res. 2009 Aug 1;69(15):6149– 6157. PMID: 19622763

14. Lin J, Tarr PT, Yang R, Rhee J, Puigserver P, Newgard CB, Spiegelman BM. PGC-1beta in the regulation of hepatic glucose and energy metabolism. J Biol Chem. 2003 Aug 15;278(33):30843–30848. PMID: 12807885

15. Deblois G, Chahrour G, Perry MC, Sylvain-Drolet G, Muller WJ, Giguère V. Transcriptional control of the ERBB2 amplicon by ERRalpha and PGC-1beta promotes mammary gland tumorigenesis. Cancer Res. 2010 Dec 15;70(24):10277–10287. PMID: 20961995

16. Chun SY, Johnson C, Washburn JG, Cruz-Correa MR, Dang DT, Dang LH. Oncogenic KRAS modulates mitochondrial metabolism in human colon cancer cells by inducing HIF-1α and HIF-2α target genes. Mol Cancer. 2010 Nov 13;9:293. PMCID: PMC2999617

17. Götzke H, Kilisch M, Martínez-Carranza M, Sograte-Idrissi S, Rajavel A, Schlichthaerle T, Engels N, Jungmann R, Stenmark P, Opazo F, Frey S. The ALFA-tag is a highly versatile tool for nanobody-based bioscience applications. Nat Commun. 2019 Sep 27;10(1):4403. PMCID: PMC6764986

18. Hernandez C, Molusky M, Li Y, Li S, Lin JD. Regulation of Hepatic ApoC3 Expression by PGC-1β Mediates Hypolipidemic Effect of Nicotinic Acid. Cell Metab. Elsevier; 2010 Oct 6;12(4):411–419. PMID: 20889132

19. Lelliott CJ, Medina-Gomez G, Petrovic N, Kis A, Feldmann HM, Bjursell M, Parker N, Curtis K, Campbell M, Hu P, Zhang D, Litwin SE, Zaha VG, Fountain KT, Boudina S, Jimenez-Linan M, Blount M, Lopez M, Meirhaeghe A, Bohlooly-Y M, Storlien L, Strömstedt M, Snaith M, Oresic M, Abel ED, Cannon B, Vidal-Puig A. Ablation of PGC-1beta results in defective mitochondrial activity, thermogenesis, hepatic function, and cardiac performance. PLoS Biol. 2006 Nov;4(11):e369. PMCID: PMC1634886

20. Victorino VJ, Barroso WA, Assunção AKM, Cury V, Jeremias IC, Petroni R, Chausse B, Ariga SK, Herrera ACSA, Panis C, Lima TM, Souza HP. PGC-1β regulates HER2-overexpressing breast cancer cells proliferation by metabolic and redox pathways. Tumour Biol J Int Soc Oncodevelopmental Biol Med. 2016 May;37(5):6035–6044. PMID: 26602383

21. Bensard CL, Wisidagama DR, Olson KA, Berg JA, Krah NM, Schell JC, Nowinski SM, Fogarty S, Bott AJ, Wei P, Dove KK, Tanner JM, Panic V, Cluntun A, Lettlova S, Earl CS, Namnath DF, Vázquez-Arreguín K, Villanueva CJ, Tantin D, Murtaugh LC, Evason KJ, Ducker GS, Thummel CS, Rutter J. Regulation of Tumor Initiation by the Mitochondrial Pyruvate Carrier. Cell Metab. 2020 Feb;31(2):284-300.e7.

22. Schell JC, Olson KA, Jiang L, Hawkins AJ, Van Vranken JG, Xie J, Egnatchik RA, Earl EG, Deberardinis RJ, Rutter J. A role for the mitochondrial pyruvate carrier as a repressor of the Warburg Effect and colon cancer cell growth. Mol Cell. 2014 Nov 6;56(3):400–413. PMCID: PMC4268416

23. Lin J, Puigserver P, Donovan J, Tarr P, Spiegelman BM. Peroxisome proliferator-activated receptor gamma coactivator 1beta (PGC-1beta), a novel PGC-1-related transcription coactivator associated with host cell factor. J Biol Chem. 2002 Jan 18;277(3):1645–1648. PMID: 11733490

