## Supplemental File 1 for "PGC-1β and ERRα promote glutamine metabolism and colorectal cancer survival via transcriptional regulation of PCK2"

pLKO-shRNA-puro sequences

| | |
|---|---|
| PGC-1β #1 | 5'CGAGCTCTCACTGCTGCAGAA |
| PGC-1β #2 | 5'GGCGGACAGCACCCAAGACAA |
| ERRα #1 | 5'GCAGAGCAATAACACTATATT |
| ERRα #2 | 5'CCCTGCAGAGCAATAACACTA |
| PCK2 #1 | 5'GTAGAGAGCAAGACGGTGATT |
| PCK2 #2 | 5'GCACATCCCAACTCTCGATTT |
