## Supplemental File 2 for "PGC-1β and ERRα promote glutamine metabolism and colorectal cancer survival via transcriptional regulation of PCK2"

5’ homology arm with 2X Strep tag and TEV site (HOM 1)

TCCCCGACCTGCAGCCCAGCTCCCAGCACTTTGGGAGGCCGAGGCAGGAAGATCATGAG
GTCAAGAGATCGAGACCATCGTGGCCAACATGGTGAAACCCCCTCTCTATTAAAAATACAA
AAATTAGCTGGGCGTGGTGGCGCACACCTGTAGTCCCAGCAACTTGGGAGGCTGAGGCG
GGAGAATCACTTGAACCCGAGAGGCAGAGGTTGCAGTGAGCTGAGATCACATCCCTGCAC
TCCAGCCTTGGTGACAGAGCGAGACTGCATCTCAAAAAAAAATAAATTCTAGTGTTGTTAG
CCCTCCCTTATGTGGCACTGCAGCAGGTTACTAATGGACGAGAAGCTGTTGGGGGAAGTA
GAGTTGTAGGGTGTTGGAGCTAGAAAGGCTCTGGAGTGCTCTAGTTTGGGCCTCCAGGTC
TCTAGATAGGACAGCCAAGGCCCTGAGACACTGATACCATGGCCAAGATGTCCCAGCAGG
ATGGCAGGCAGTGCCCACAGTCCAGCGCTCATCCCAGCTCCCCAAGGCCTTGGGCTGCA
GCCACTCCACGGTGCCCACTCATAGTGGCAGATTCTTCAACCATCCGGTCTTTGTAAGTTG
CTCACTGCCTTCCCCTCTTCCCTGCCTCTTCACCCCCATGCCCAGATTCCAATTCAGAAGA
GGCCCTTCCTGCGTCAGGGAAAAGCAAGTATGAAGCCATGGATTTTGACAGCTTACTGAAA
GAGGCCCAGCAGAGCCTGCATGAGAACCTGTACTTCCAATCCAATGGGAGCGGAGGAGG
TTCCGG

3’ homology arm (HOM 2)

AGTTCTTCTGATTCGAACATCAGACAAGGCCCTTCCAATATGTTTACGTTTTCAAAGAAATC
AAGTATATGAGGAGAGCGAGCGAGCGTGAGAGAACACCCGTGAGAGAGACTTGAAACTGC
TGTCCTTTAAAAAAAAAAAAAATCAATGTTTACATTGAACAAAGCTGCTTCTGTCTGTGAGTT
TCCATGGTGTTGACGTTCCACTGCCACATTAGTGTCCTCGCTTCCAACGGGTTGTCCCGGG
TGCACCTCGAAGTGCCGGGTCCGTCACCCATCGCCCCTTCCTTCCCGACTGACTTCCTCT
CGTAGACTTGCAGCTGTGTTCACCATAACATTTCTTGTCTGTAGTGTGTGATGATGAAATTG
TTACTTGTGAATAGAATCAGGACTATAAACTTCATTTTTAATTGAAAAAAAAAGTATATCCTT
AAAATAATGTATTTATGGCTCAGATGTACTGTGCCTGGGATTATTGTATTGCTTCCTTGATTT
TTTAACTATGCACTGTCATGAGGTGTTTGCCACTGAGCTGCCCTGCTCCCCTTGCCAGATT
GCCCTGGAGGTGCTGGGTGGCCGCTAGGCTGGTCTGCAGGAAAGCGCGGCCTGCCGTTT
CCGGGCCGTATCTGCCAAGCCCTGCCTTGTCTCTTACTGAGCAAGTTTGGCTCAAATTATA
GGAGCCCCCATCTTGTGCCCAGCTCATGCTCCAAGTGTGTGTCTATCCATTTGTACTCAGA
CTCTTGAGTACCTTGTAAGGAAGGCGGGGCAAGCTCTTGGAAAGTCCTCTCCA

Cas 9 gRNA sequence to cut PGC-1β adjacent to the stop codon:

5’ GATAACAGCCTTAACCCTCG

PCR primers for verification of genomic insertion

Forward primer 5’ CAGGACTGAGTGTCAGTGACC

Reverse primer 5’ CAGCAGGCTGAAGTTAGTAGC
