## Supplemental File 3 for "PGC-1β and ERRα promote glutamine metabolism and colorectal cancer survival via transcriptional regulation of PCK2"

LLxxL 1 into AAxxA

```
            E   N   E   A   A   A   A   E   A   T   K   T   L
LA1-F 5'GT GAG AAT GAG GCC GCC GCT GCA GAG GCC ACC AAG ACC CTG G
LA1-R 5'C CAG GGT CTT GGT GGC CTC TGC AGC GGC GGC CTC ATT CTC AG
```

LxxLL 2 into AxxAA

```
           E   L   S   L   A   Q   K   A   A   L   A   T   S   Y
LA2-F 5'C GAG CTC TCA CTG GCT CAG AAG GCC GCC CTG GCC ACA TCC TAC
LA2-R 5'GTA GGA TGT GGC CAG GGC GGC CTT CTG AGC CAG TGA GAG CTC G
```

LxxLL 3 into AxxAA

```
         E   F   S   I   A   R   E   A   A   A   Q   D   V   L
LA3-F 5'GAG TTC TCC ATT GCC AGG GAA GCT GCT GCT CAA GAC GTG CTC
LA3-R 5'GAG CAC GTC TTG AGC AGC AGC TTC CCT GGC AAT GGA GAA CTC
```

LLxxL 4 into AAxxA

```
            E   R   S   E   A   A   S   H   A   R   H   A   T
LA4-F 5'CA GAG CGA AGT GAG GCC GCT TCC CAC GCC CGA CAT GCC ACA G
LA4-R 5'C TGT GGC ATG TCG GGC GTG GGA AGC GGC CTC ACT TCG CTC TG
```

LRELL into LAALL (RAEA)

```
             E   F   S   I   L   A   A   L   L   A   Q   D   V
RAEA-F 5'CT GAG TTC TCC ATT CTG GCC GCC CTT CTG GCT CAA GAC GTG
RAEA-R 5'CAC GTC TTG AGC CAG AAG GGC GGC CAG AAT GGA GAA CTC AG
```
