## Supplemental File 4 for "PGC-1β and ERRα promote glutamine metabolism and colorectal cancer survival via transcriptional regulation of PCK2"

siRNA sequences and product number

| Target | Item # | Target Sequence |
|---|---|---|
| PGC1$\beta$ | J-008556-06 | CCAGAAGGCGUCCUGCAAA |
| PGC1$\beta$ | D-008556-02 | GUACAGAACUACAUAAGCA |
| ERR$\alpha$ | J-003403-07 | GGCCUUCGCUGAGGACUUA |
| ERR$\alpha$ | J-003403-08 | GCGAGAGGAGUAUGUUCUA |
| PCK2 | J-006797-07 | GAGCAAGACGGUGAUUGUA |
| PCK2 | J-006797-09 | CCUGGGAGAUGGUGACUUU |
| Non-Targeting 1 | D-001810-01 | UGGUUUACAUGUCGACUAA |
| Non-Targeting 2 | D-001810-02 | UGGUUUACAUGUUGUGUGA |
