## Supplemental File 5 for "PGC-1β and ERRα promote glutamine metabolism and colorectal cancer survival via transcriptional regulation of PCK2"

| siRNA PGC1$\beta$ | | | siRNA ERR$\alpha$ | | |
|---|---|---|---|---|---|
| **Gene** | **Fold Change ($Log_2$)** | **Adjusted P-Value** | **Gene** | **Fold Change ($Log_2$)** | **Adjusted P-Value** |
| ESRRA | -3.0501725 | 4.34E-23 | SLC1A4 | -3.8956646 | 2.63E-12 |
| PCK2 | -2.9644106 | 2.35E-13 | PCK2 | -3.9065517 | 5.96E-12 |
| SLC1A4 | -2.7414698 | 2.65E-12 | JDP2 | -2.8422453 | 2.00E-10 |
| TRIB3 | -1.82452 | 1.82E-08 | LMNB1 | -2.6485427 | 2.40E-10 |
| PSAT1 | -1.6115383 | 2.54E-08 | CHAC1 | -2.91747 | 6.56E-08 |
| DDIT4 | -2.0444729 | 4.67E-08 | C1orf116 | 2.80287776 | 7.74E-08 |
| CPA4 | 1.76323906 | 7.60E-08 | HR | -2.0447284 | 4.57E-07 |
| GPT2 | -1.6992819 | 1.01E-07 | MDK | -1.7412193 | 1.17E-06 |
| ULBP1 | -1.9837393 | 2.01E-07 | NRP1 | 2.87549936 | 4.34E-06 |
| ASNS | -2.4863819 | 4.64E-07 | TFPI | 1.87366954 | 4.55E-06 |
| JDP2 | -1.9716087 | 7.61E-07 | PHGDH | -2.3778323 | 5.27E-06 |
| IL6R | 1.48849468 | 1.09E-06 | FHL1 | -2.6007623 | 9.10E-06 |
| SEMA7A | 1.73161535 | 1.17E-06 | ASNS | -2.2820422 | 2.43E-05 |
| KRT80 | 2.30875312 | 1.00E-05 | CPA4 | 2.25251952 | 4.07E-05 |
| NR6A1 | 1.70161587 | 1.18E-05 | ZBTB20 | 2.25834811 | 8.26E-05 |
| BIRC3 | 2.1938021 | 2.89E-05 | CEACAM1 | 3.14424259 | 8.58E-05 |
| SLC43A1 | -2.8855272 | 3.52E-05 | LMO7 | 2.19563308 | 8.58E-05 |
| AMMECR1 | 1.19063512 | 3.58E-05 | PALM3 | -3.4196186 | 0.00010345 |
| PDGFB | 1.7233171 | 3.98E-05 | AMPD3 | 1.65302827 | 0.00010467 |
| LRATD2 | -1.2124766 | 4.54E-05 | SNAI2 | 3.16959983 | 0.0001191 |
| PHGDH | -1.4993388 | 0.00013067 | PSIP1 | -1.7414462 | 0.0001191 |
| STC2 | -1.150869 | 0.00015699 | BTG2 | 1.95825153 | 0.0001312 |
| KCNH3 | -1.6289723 | 0.00015699 | KDM5B | 1.50327189 | 0.00013924 |
| NACC2 | 1.08933939 | 0.00015699 | NUP210 | -1.841356 | 0.00013924 |
| SH3RF2 | 1.81821689 | 0.00036751 | DOCK4 | 1.72009796 | 0.00014812 |
| SOCS1 | 2.24381519 | 0.00036751 | EIF5A2 | -1.8039583 | 0.00014812 |
| ANKRD52 | 1.32571374 | 0.00037372 | COL17A1 | 2.60487877 | 0.0001486 |
| PDCD4 | -1.5262688 | 0.00053679 | GPT2 | -1.7892692 | 0.00014877 |
| KRT23 | -1.5967489 | 0.00061723 | ZNF367 | -2.5428911 | 0.00015907 |
| ESRP1 | -0.9707887 | 0.00068048 | THEM6 | -1.7297649 | 0.00017903 |

Genes with significant differential expression following depletion of PGC1$\beta$ or ERR$\alpha$ compared to a non-targeting control.
